# The folding and unfolding behavior of ribonuclease H on the ribosome

**DOI:** 10.1101/2020.04.16.044867

**Authors:** Madeleine K. Jensen, Avi J. Samelson, Annette Steward, Jane Clarke, Susan Marqusee

## Abstract

The health of a cell depends on accurate translation and proper protein folding; misfolding can lead to aggregation and disease. The first opportunity for a protein to fold occurs during translation, when the ribosome and surrounding environment can affect the energy landscape of the nascent chain. However, quantifying these environmental effects is challenging due to the ribosomal proteins and rRNA, which preclude most spectroscopic measurements of protein energetics. We have applied two gel-based approaches, pulse proteolysis and force-peptide arrest assays, to probe the folding and unfolding pathways of RNase H ribosome-stalled nascent chains. We find that ribosome-stalled RNase H has an increased unfolding rate compared to free RNase H, which completely accounts for observed changes in protein stability and indicates that the folding rate is unchanged. Using arrest peptide-based force-profile analysis, we assayed the force generated during the folding of RNase H on the ribosome. Surprisingly, we find that population of the RNase H folding intermediate is required to generate sufficient force to release the SecM stall and that readthrough of the stall sequence directly correlates with the stability of the folding intermediate. Together, these data imply that the folding pathway of RNase H is unchanged on the ribosome. Furthermore, our data indicate that the ribosome promotes unfolding while the nascent chain is proximal to the ribosome, which may limit the deleterious effects of misfolding and assist in folding fidelity.

## INTRODUCTION

In the cell, a protein has its first opportunity to fold during synthesis on the ribosome. It is critical, therefore, that this initial folding event proceeds with fidelity and avoids toxic misfolded states (1, 2). Both the vectorial nature of translation and interactions with the ribosome can affect this process (3–8), such that the co-translational folding pathway can be different from the pathway observed during re-folding experiments. For instance, co-translational folding of firefly luciferase promotes formation of an intermediate that helps to prevent misfolding (9). For the protein HaloTag, co-translational folding avoids an aggregation-prone intermediate and leads to more efficient folding (10). Both of these proteins have complex folding pathways, including the formation of transient intermediates, and both show differences between their co-translational and refolding pathways. However, folding pathways are not necessarily altered by the ribosome; two small beta-sheet domains, the src SH3 domain and titin I27, show simple two-state folding and appear to fold via the same pathway both on and off the ribosome (5, 11). Is populating transient folding intermediates required for folding to be modulated by the ribosome? To understand how proteins fold *in vivo*, it is essential to elucidate how the ribosome affects nascent chain folding.

Measuring the energetics and dynamics of ribosome-stalled nascent chains (RNCs) presents numerous experimental challenges. Historically, the kinetics of protein folding have been monitored with spectroscopic techniques, such as circular dichroism (CD) or fluorescence; structural features of the folding trajectory can be further probed by hydrogen-deuterium exchange (HDX) and by comparing stabilities and folding rates among site-specific variants of the protein of interest (ϕ-value analysis) (12). These well-established techniques, however, are not suitable for monitoring folding of an RNC; the ribosome contains over 50 proteins, which makes it impossible to specifically detect and measure folding of the nascent chain by most spectroscopic methods. While more complex biophysical approaches, such as NMR (4, 6), Förster resonance energy transfer and photoinduced electron transfer (13, 14), and optical trap mechanical studies (3, 11, 15, 16), have elucidated important features of the folding process recently, these approaches are very technically challenging and have significant throughput limits. Thus, it is hard to know if findings using these techniques are generally applicable to the entire proteome.

An alternative technique to measure folding kinetics and energetics in a complex mixture is pulse proteolysis. Under appropriate conditions, a short pulse of proteolysis degrades unfolded proteins, leaving the population of folded proteins intact. Unlike limited proteolysis, which reports on the relative proteolytic susceptibility of a protein, pulse proteolysis can be used to measure the fraction of folded protein in a given mixture quantitatively. Pulse proteolysis has been applied under equilibrium conditions to determine protein stability (ΔG_unf_) or in a kinetic experiment to assess unfolding kinetics (*k*_unf_) and requires very little protein relative to spectroscopic approaches (17–19). Since the fraction of folded protein is quantified by following a specific band on the gel, it can be carried out in complex mixtures, making it ideally suited for the study of RNCs.

We recently used pulse proteolysis to monitor the thermodynamic stability of proteins off and on the ribosome with varying linker lengths (or distances) from the peptidyl-transferase center (PTC) (20). For all proteins studied, the ribosome destabilizes nascent chains compared to free protein, and this destabilization is dependent on the distance from the PTC (20). The physical factors behind this destabilization remain unclear. Is the destabilization rooted in the folding and/or unfolding kinetics of the nascent chains? Pulse proteolysis applied in a kinetic mode can assess how the ribosome affects nascent chain folding and unfolding.

Another approach recently developed to explore the folding of RNCs is arrest peptide-based force-profile analysis (FPA) (21). FPA monitors the release of a stalled nascent chain, which has been shown to correlate with folding near the exit tunnel (5, 21–23), and has also been shown to be related to the global stability and folding topology of the nascent chain (24, 25). Despite several publications using this technique, details about the types of folding events required to release a stall are unclear. Release is usually attributed to global folding; however, force profiles of HemK and DHFR suggest that formation of an intermediate structure can also trigger release, resulting in bimodal or multi-modal force profiles (23, 26).

Here, we apply pulse proteolysis to determine the unfolding rate of RNase H RNCs and combine this technique with FPA to understand how the ribosome modulates the folding and unfolding of RNase H (RNH*, * denotes a cysteine-free variant) and its variants. For ribosome-stalled RNH* I53D (a two-state folding variant of RNH*), the thermodynamic destabilization can be attributed to an increase in the unfolding rate, indicating that the folding rate is unaltered on the ribosome. Force-profile studies on two-state and three-state folding variants of RNH* show that both the presence and stability of the folding intermediate are critical for force generation and readthrough of the stall sequence. Together these results suggest that the folding pathway for RNH* is the same on and off the ribosome and indicate that folding to a stable native state does not necessarily cause arrest-peptide release, while formation of a transiently-populated intermediate is sufficient for release. These results have implications for understanding the fundamental principles of both co-translational folding and membrane protein translocation.

## RESULTS

### Stability and kinetics of RNase H monitored by CD

The protein RNase H* (RNH*) has been extensively characterized in bulk by both CD and tryptophan fluorescence. The detail with which we understand its energy landscape make it a prime candidate for further studies on the ribosome (27, 28). These data, however, were all obtained in a buffered solution at pH 5.5, not the pH 7.4 buffer conditions (with divalent cations) required for on-the-ribosome studies. Therefore, we measured the stability and unfolding kinetics of RNH* I53D by CD in a buffer that both more closely approximates physiological conditions and is suitable for ribosome-bound experiments (pH 7.4, 150 mM KCl, 15 mM Mg(OAc)_2_, 0.1 mM TCEP). Compared to the previous studies at pH 5.5, RNH* I53D is slightly destabilized (ΔG_unf_ = 4.71 ± 0.32 kcal/mol versus 5.6 ± 0.4 kcal/mol) (Figure 1 and Table 1). The extrapolated unfolding rate in the absence of denaturant is slightly higher [2.0 (± 0.5) × 10^−5^ s^-1^ at pH 7.4 compared to 6.3 (± 5) × 10^−6^ s^-1^ at pH 5.5] (Figure 1 and Table 1), and *m*^‡^ _unf_ = 0.45 ± 0.02 kcal mol^-1^ M^-1^ at pH 7.4, within error of the *m*^‡^ _unf_ measured previously for RNH* I53D (0.5 ± 0.1 kcal mol^-1^ M^-1^, pH 5.5) (21). These data serve as a point of comparison for the gel-based studies comparing folding on and off the ribosome.

**Table 1.** Stability and unfolding kinetics of RNH* I53D. ^a^ Values from Samelson *et al*. (20). ^b^ Calculated using *m*_unf_ from CD.

|  | RNH* I53D (CD) | RNH* I53D-(GS)<sub>5</sub>-<br>SecM (off) | RNH* I53D-(GS)<sub>5</sub>-<br>SecM (on) |
| --- | --- | --- | --- |
| $C_m$ (M, urea) | $2.11 \pm 0.09$ | $2.14 \pm 0.13^a$ | $1.65 \pm 0.10^a$ |
| $\Delta G_{\text{unf}}$ (kcal/mol) | $4.71 \pm 0.32$ | $4.77 \pm 0.29^b$ | $3.68 \pm 0.22^b$ |
| $m_{\text{unf}}$ (kcal mol<sup>-1</sup> M<sup>-1</sup>) | $2.23 \pm 0.06$ | - | - |
| $k_{\text{unf}}^{\text{0M urea}}$ (s<sup>-1</sup>) | $2.0 (\pm 0.5) \times 10^{-5}$ | $4.1 (\pm 2.4) \times 10^{-5}$ | $3.9 (\pm 3.7) \times 10^{-4}$ |
| $m_{\text{unf}}^{\ddagger}$ (kcal mol<sup>-1</sup> M<sup>-1</sup>) | $0.45 \pm 0.02$ | $0.35 \pm 0.08$ | $0.40 \pm 0.94$ |

**Figure 1.**
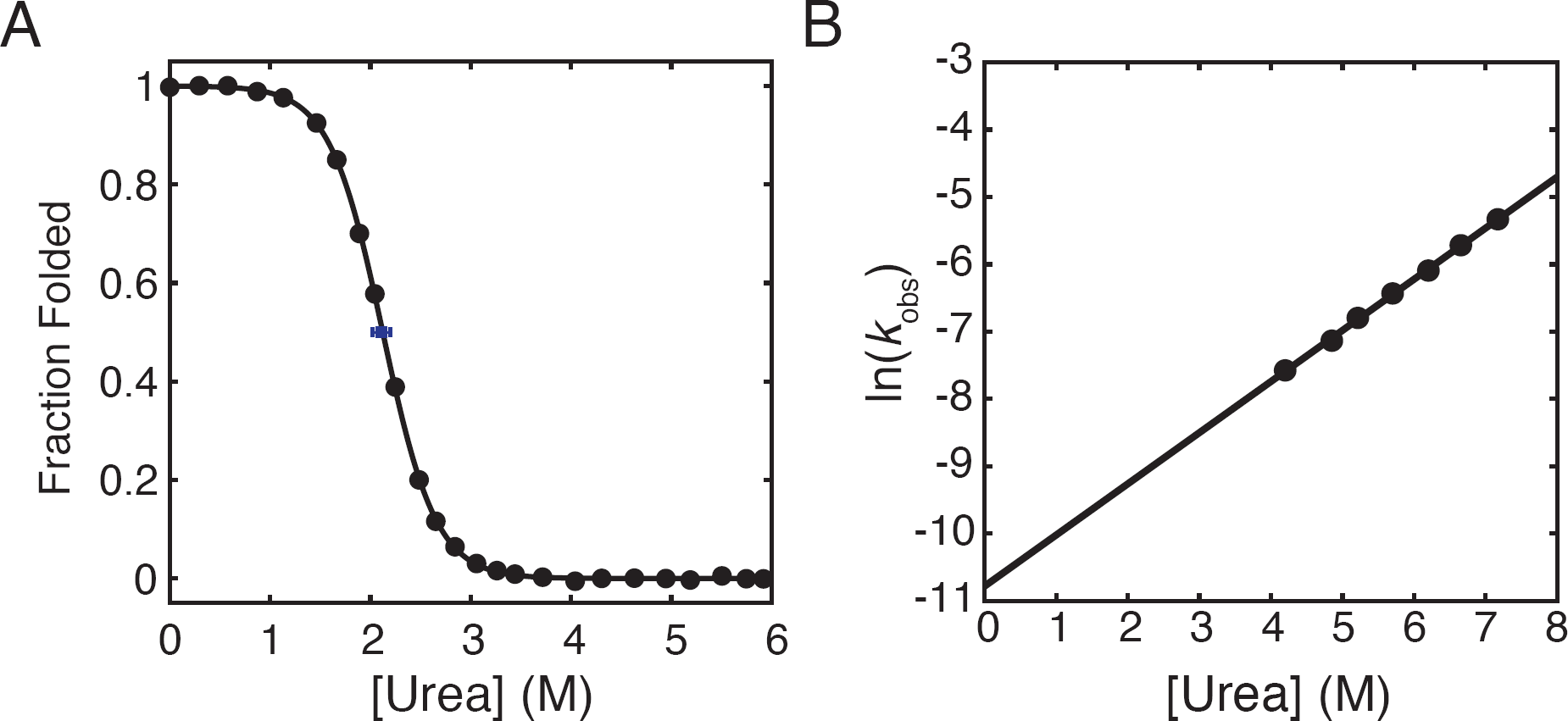
Stability and unfolding kinetics of RNH* I53D measured by CD at pH 7.4. A) Equilibrium urea-induced denaturant melt (black circles) of RNH* I53D (pH 7.4 at 25 °C). Data were fit using a two-state approximation (U⇋N), and the data shown are from one representative experiment of three. The average C_m_ from three experiments is marked by a blue square with error bars for the standard deviation (SD). B) Natural log of the observed unfolding rate for RNH* I53D at pH 7.4 as a function of urea.

### Ribosome-tethered RNH* I53D has a higher unfolding rate than free protein

Previous pulse proteolysis studies determined the equilibrium stability (ΔG_unf_) of RNH* I53D on and off the ribosome (20). The observed destabilization on the ribosome implies an underlying change in the folding and/or unfolding kinetics. If the two-state folding mechanism of RNH* I53D holds true as an RNC, measurements of the unfolding rate via pulse proteolysis will allow us to infer changes in the folding rate (K_eq_=*k*_unf_/*k*_fold_).

Unfolding kinetics were monitored using pulse proteolysis by rapidly diluting a protein or RNC sample to a specific final urea concentration and following the fraction folded as a function of time by assaying with a pulse of thermolysin-based proteolysis (see Methods). Our pulse length of one minute allows us to measure unfolding rates on the order of ∼10^−2^ sec^-1^ or less, which partially dictates the urea concentrations accessible for these measurements. For studies on RNCs, experiments are also limited to below 3.5 M urea to maintain the integrity of the ribosome, which dissociates above 3.5 M urea (20).

Nascent chains with a C-terminal ten-residue glycine-serine (GS) linker and the well-characterized SecM stalling sequence were tagged by incorporating BODIPY-FL-Lysine^AAA^-tRNA (Promega) during IVT (20) (see Figure 2A and Methods). Off the ribosome, RNH* I53D-(GS)_5_-SecM unfolds with *k*_unf_^0M urea^ = 4.1 (± 2.4) × 10^−5^ s^-1^ and *m*^‡^_unf_ = 0.35 ± 0.08 kcal mol^-1^ M^-1^ (Figure 2B and D), within error of that determined by CD (Table 1). On the ribosome, we measured *k*_unf_^0M urea^ to be 3.9 (± 3.7) × 10^−4^ s^-1^, an order of magnitude greater than off the ribosome (Figures 2C and D and Table 1). The *m*^‡^-values are comparable on and off the ribosome, with *m*^‡^_unf_ = 0.40 ± 0.94 kcal mol^-1^ M^-1^ for the RNC (Figure 2D and Table 1). Assuming that RNH* I53D is two-state on the ribosome, *k*_fold_ = 0.2 s^-1^, similar to that obtained by pulse proteolysis off the ribosome (*k*_fold_ = 0.1 s^-1^, pH 7.4) and from bulk CD experiments (0.1 s^-1^, pH 5.5) (27).

**Figure 2.**
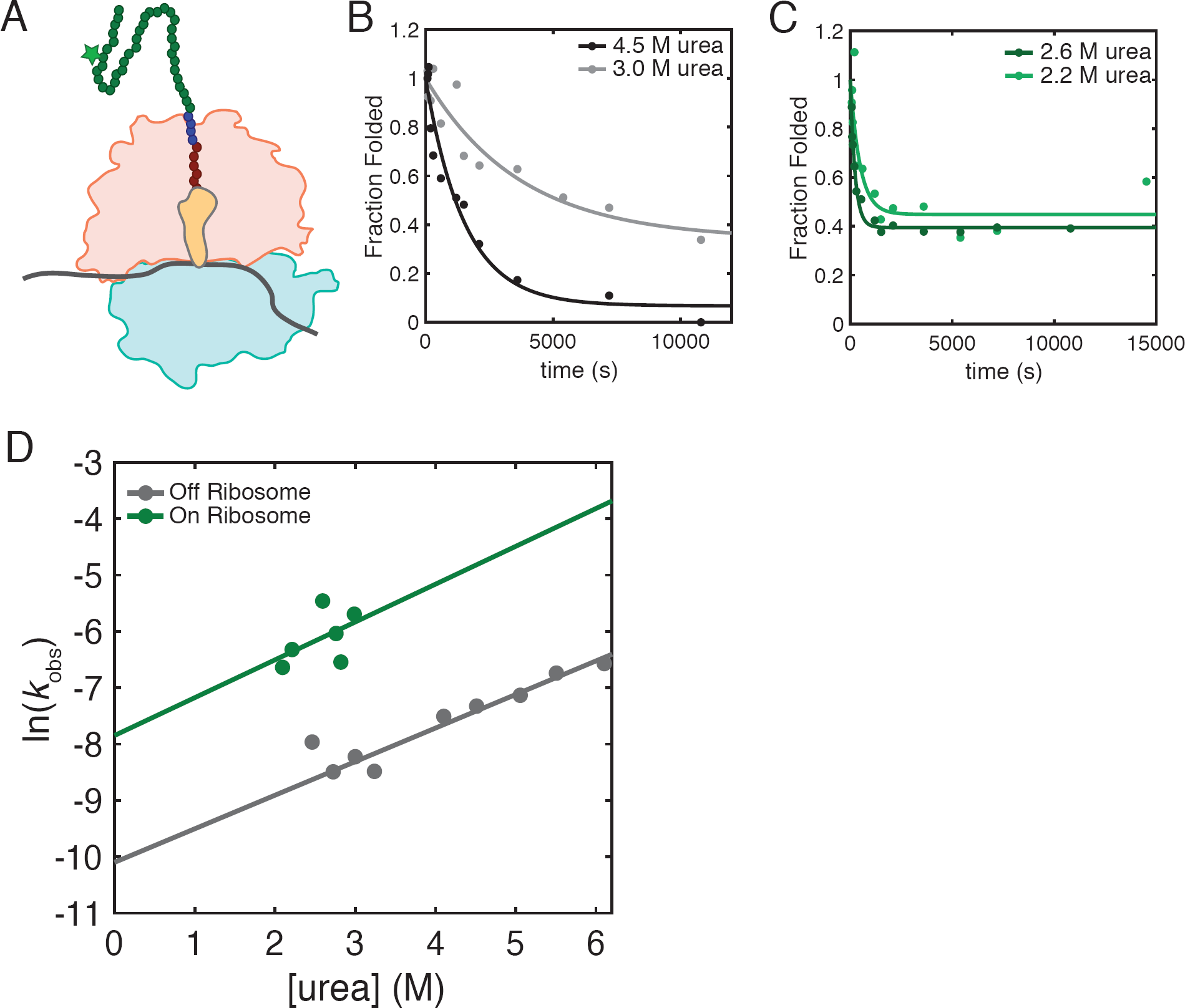
Unfolding kinetics as measured by pulse proteolysis. A) Cartoon representation of a ribosome-stalled nascent chain generated for pulse proteolysis experiments. A 50S subunit (orange) and 30S subunit (light blue) with a peptidyl-tRNA (yellow) stalls during translation of mRNA (grey) at a SecM stall sequence (red circles). The protein of interest (green circles) is extended beyond the ribosome exit tunnel by a ten-residue glycine-serine linker (blue circles). The nascent chain is tagged with a BODIPY-FL-Lysine (star). B) Representative traces of observed off-ribosome unfolding rates of RNH* I53D at 4.5 M urea (black) and 3.0 M urea (grey). C) Representative traces of observed on-ribosome unfolding rates of RNH* I53D at 2.6 M urea (dark green) and 2.2 M urea (light green). D) Data sets including (B) and (C) are fit to a single exponential to extract *k*_obs_, and the ln(*k*_obs_) for experiments on (green) and off the ribosome (grey) are plotted at several urea concentrations. These data are fit to a linear model where the slope is *m*^‡^ _unf_ and the y-intercept is *k* _unf_^0M urea^, the unfolding rate in a 0 M denaturant condition. The urea concentrations at which we can investigate the unfolding rates of nascent chains are limited based on the C_m_ of the nascent chain and the stability of 70S ribosomes, leading to a higher error for the extrapolation of on-ribosome data.

### RNH* I53D does not read through the SecM stall

The folding trajectory of ribosome-bound nascent chains can be interrogated directly using arrest peptide-based force profile analysis (23). Arrest peptides, such as the SecM stalling sequence, are highly sensitive to tension on the nascent chain, and tension generated due to folding has been shown to release arrest (5, 21–23, 26). Force-profile analysis (FPA) measures the fraction readthrough of the SecM stall (*f*_FL_) as a function of linker length between the protein of interest and the PTC. At each linker length, the *f*_FL_ is a readout of the fraction of nascent chains that release the stall. The connection between the biophysical properties of the nascent chain and the *f*_FL_ remains unclear. The stability of the nascent chain, its topology, and folding rate have all been linked to the amplitude of the *f*_FL_ (24, 25). Thus, a comparison of FPA on the well-characterized protein RNH* with the on-ribosome energetics and kinetics obtained by pulse proteolysis should help to decipher these effects, in addition to reporting on the RNH* folding trajectory on the ribosome.

Figure 3A shows the force profiles of both RNH* I53D and a non-folding control. Four mutations were needed to generate the non-folding control (F8A/I25A/I53D/W85A) based on data derived from bulk ensemble studies on variants of RNH* and assumed additivity (27, 29, 30). As expected, the non-folding variant shows minimal release of the stall sequence across all linker lengths tested (Figure 3A). Surprisingly, the force-profile assay for RNH* I53D resembles the non-folding variant (Figure 3A). This result is not due to a lack of the ability of RNH* I53D to fold on the ribosome, as pulse proteolysis data confirm that RNH* I53D can fold on the ribosome with a 35-residue linker (20). Importantly, the calculated on-ribosome *k*_fold_ indicates that RNH* I53D-(GS)_5_-SecM has ample time to fold during FPA, where samples are incubated for 15 min prior to measuring release (see above). In spite of this, folding does not appear to generate the force needed for release of the SecM-mediated stall.

**Figure 3.**
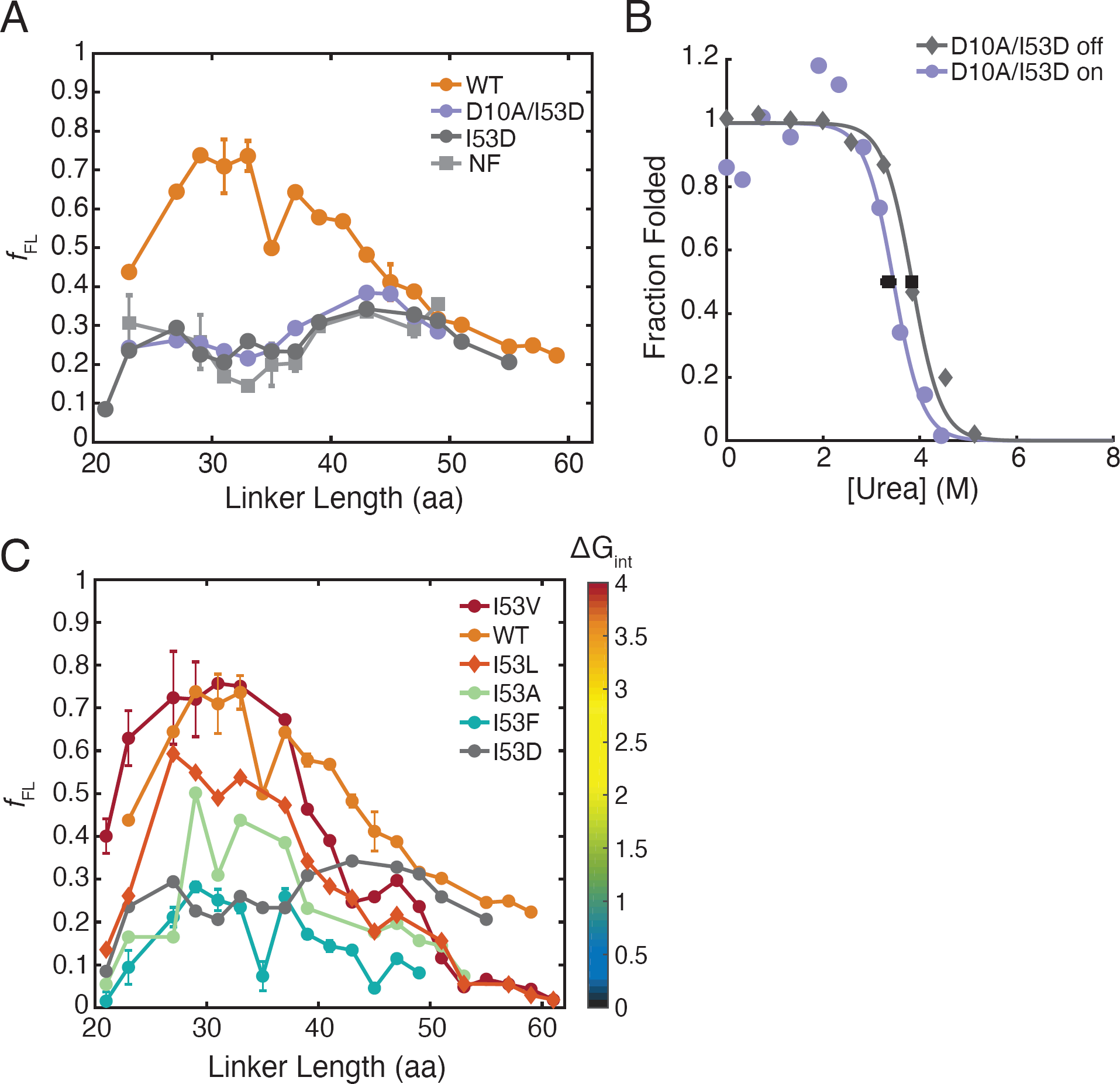
Force-profile analysis of RNH* variants. A) FPA of RNH* wildtype (orange circles), a non-folding control (NF, light grey squares), which is RNH* F8A/I25A/I53D/W85A, RNH* I53D (dark grey circles), and RNH* D10A/I53D (purple circles). The lines connect the data to show trends and are not fits. Where L## denotes a linker length, the following are the average of two experiments: WT L23, L27, L31, L33, L37, L39, L43, L45; NF L43. NF L23, L27, L37, L39, and L47 are the average of three experiments. All other points shown are the result of a single experiment. Error bars shown are standard deviations. B) Stability of RNH* D10A/I53D-(GS)_5_-SecM measured off (grey) and on (purple) the ribosome by pulse proteolysis from IVT-produced (PURExpress ΔRF123) protein. Data shown are representative of three experiments off and two experiments on the ribosome with black squares marking the average C_m_s from on and off the ribosome samples and error bars showing the standard deviations. Data are fit using a two-state approximation. C) FPA of RNH* I53 variants: I53V (red circles), I53L (orange diamonds), I53A (green circles), and I53F (blue circles). WT (orange circles) and RNH* I53D (grey circles) are repeated from (A) for reference. The heat map shows the color coding of the variants based on the stability of their folding intermediate (ΔG_int_). Lines connect the data to show trends and are not fits. I53V L21, L23, L27, and L29, as well as I53F L21, L23, L27, L29, L31, L33, L35, L37, L39, and L41 are the average of two experiments with the standard deviation shown as error bars. All other points are the result of single experiments.

### Three-state RNH* can read through SecM stall

To determine if this uncoupling of folding and SecM readthrough holds for other RNases H, we then turned to RNH*, which is known to fold in a three-state manner (U⇌I⇌N) both in bulk and in single-molecule optical trap experiments (27, 31). In bulk ensemble studies, RNH* is also more stable than RNH* I53D (ΔG_unf_ = 9.7 versus 5.6 kcal/mol, pH 5.5) (27, 32). Figure 3A shows that the force profile of RNH* is notably different than those observed for RNH* I53D and the non-folding control. Unlike the others, RNH* shows significant release (*f*_FL_ > 0.5) occurring at linker lengths from 29 to 41 residues. Thus, RNH* folding is capable of generating the force necessary to release the stall.

### Increased global stability does not result in SecM readthrough

We made a stabilizing variant of RNH I53D, RNH* D10A/I53D, which has a stability near that of wildtype RNH* (ΔG_unf_ = 8.5 kcal/mol, pH 5.5) and, like RNH* I53D, shows two-state folding in bulk studies (30). This construct would allow us distinguish whether SecM readthrough is failing due to stability or population of the intermediate. The off-ribosome stability of RNH* D10A/I53D-(GS)_5_-SecM is 7.66 ± 0.03 kcal/mol versus 6.71 ± 0.28 kcal/mol as a stalled nascent chain. Both are significantly more stable than RNH* I53D (Figure 3B and Table 2). Despite its increased stability, this variant also failed to release the stall at all linker lengths (Figure 3A). Thus, the release observed for RNH* is not solely due to its high stability. This suggests that it is either the formation of the RNase H folding intermediate or the process of folding from the intermediate that is responsible for force generation on the SecM stall sequence.

**Table 2.** Stability of RNH* D10A/I53D. ^a^ Values from Connell *et al*. (30). Off-ribosome values are from three experiments and on-ribosome values are from two experiments.

|  | RNH* D10A/I53D<br>(CD)<sup>a</sup> | RNH* D10A/I53D-(GS)<sub>5</sub>-<br>SecM (off) | RNH* D10A/I53D-(GS)<sub>5</sub>-<br>SecM (on) |
| --- | --- | --- | --- |
| $C_m$ (M, urea) | $4.25 \pm 0.2$ | $3.83 \pm 0.02$ | $3.35 \pm 0.14$ |
| $\Delta G_{\text{unf}}$ (kcal/mol) | $8.5 \pm 0.4$ | $7.66 \pm 0.03$ | $6.71 \pm 0.28$ |
| $m_{\text{unf}}$ (kcal mol<sup>-1</sup> M<sup>-1</sup>) | $2.0 \pm 0.1$ | - | - |

### Stability of the folding intermediate correlates with arrest peptide release

To test the role of the kinetic folding intermediate in force generation and release of the stall, we turned to a series of site-specific variants known to modulate the stability of the intermediate. The folding intermediate of RNH* has been well-characterized by pulse-labeling HDX and protein engineering (30, 33, 34). Residue 53 resides in the center of helix A, an important structural feature of the intermediate. Altering the hydrophobicity and size of residue 53 is known to modulate both the stability of the intermediate (ΔG_int_) and global stability of the protein (ΔG_unf_) (27). These site-specific variants range from near-wildtype ΔG_int_ (RNH* I53V and RNH* I53L) to no detectable intermediate (two-state folding) for RNH* I53D (Table 3). To examine the effect of these mutations on the force profile of RNH*, we generated four additional substitutions (V, L, A, and F) at residue 53 for 20 linker lengths ranging from 21 to 61 residues.

**Table 3.** Kinetic folding parameters for RNH* I53 variants measured in Spudich *et al*. (27).

|  | RNH* | RNH* I53V | RNH* I53L | RNH* I53A | RNH* I53F |
| --- | --- | --- | --- | --- | --- |
| $\Delta G_{\text{unf}}$ (kcal/mol) | 9.9 | 10.6 | 9.0 | 8 | 6.3 |
| $m_{\text{unf}}$ (kcal mol<sup>-1</sup> M<sup>-1</sup>) | 2.1 | 2.3 | 2.3 | 2.4 | 2.2 |
| $\Delta G_{\text{int}}$ (kcal/mol) | 3.5 | 4.0 | 3.6 | 1.7 | 0.8 |

The stability of the intermediate appears to play the dominant role in determining the force profile. For all five variants at position 53, a higher ΔG_int_ leads to more robust readthrough of the SecM stalling sequence. RNH* I53V and RNH* I53L, which have near-wildtype ΔG_int_, have wildtype-like force profiles in terms of the linker lengths at which significant amounts of full-length protein are produced and the overall broadness of the peak (Figure 3C). RNH* I53A, for which ΔG_int_ = 1.7 kcal/mol at pH 5.5 (27, 32), did not generate more than 50% *f*_FL_ at any linker length despite its high global stability (Figure 3C). Likewise, RNH* I53F barely populates the intermediate (ΔG_int_ = 0.8 kcal/mol) (21), and readthrough of SecM was not notably different from the force profile of RNH* I53D.

## DISCUSSION

We have used a combination of pulse proteolysis and arrest peptide-based force-profile experiments to investigate how the ribosome modulates folding and unfolding of the small protein RNase H. For RNH* I53D-(GS)_5_-SecM, we observed an increase in the unfolding rate on the ribosome of about an order of magnitude compared to off the ribosome at all urea concentrations tested. This accounts for the observed decrease in stability for these RNCs. Although we can only sample a limited range of urea concentrations in these RNC experiments, the similarity in urea dependence (*m*^‡^_unf_) indicates the protein is traversing the same transition state barrier on and off the ribosome, consistent with observations for the small proteins titin and SH3 (5, 11). Assuming that RNH* I53D folds in a two-state mechanism on the ribosome (populating only U and N), this implies that the presence of the ribosome does not affect the folding rate and suggests a mechanism by which the ribosome destabilizes the nascent chain by promoting its unfolding. In general, such increases in the unfolding rate could provide a mechanism for delayed folding of the emerging nascent chain until it has extended far enough from the PTC to avoid non-native, toxic states.

In spite of its robust folding as an RNC, our studies suggest that the folding of RNH* I53D does not generate enough force to read through the SecM stall in FPA. Our pulse proteolysis studies allow us to measure both the stability and unfolding rate for RNH* I53D-(GS)_5_-SecM, indicating that RNH* I53D is capable of folding on the ribosome with a linker of 35 amino acids from the PTC. However, in our force-profile assays RNH* I53D does not read through the SecM stall at any linker length from 21 to 61 residues. Therefore, stable folding cannot be the only requirement for force generation in the force-profile assays. Moreover, increasing global stability for this protein, such as in RNH* D10A/I53D, also does not result in an increase in *f*_FL_. Previous studies have interpreted the release of translation arrest during FPA as nascent chain folding and, inversely, lack of release as either a lack of folding or folding occurring far from the ribosome surface (5, 21–23). Our results demonstrate that stable folding alone does not trigger SecM readthrough.

However, several other variants of RNH* do show appreciable force-generated release, all of which are known to populate an early intermediate in the refolding trajectory. Wildtype RNH* releases and reads through the SecM stall for a broad range of linker lengths. Additionally, by studying a series of site-specific variants, we find that ΔG_int_ is directly related to the *f*_FL_ produced. Compared to RNH*, the *f*_FL_ diminishes with decreasing ΔG_int_ for these variants, showing that we can tune the release of RNH* by adjusting the stability of the intermediate. In agreement with previous studies, the amplitude of *f*_FL_ for these three-state variants correlates with global stability (24, 25). Since the stability of the intermediate is related to the height of the subsequent folding barrier, this kinetic barrier may play a role. Interestingly, we observe release of three-state RNH* variants at lower linker lengths than would be expected based solely on the relationship between protein length and the linker length of maximum *f*_FL_ (24). Perhaps the topology of RNH* is important for this release at low linker lengths (25). Additionally, the discrepancy could be explained by release being initiated by folding of the intermediate, which contains 79 residues as compared to the 155 residues of the full-length protein. Together, our data suggest that a baseline global stability and population of the folding intermediate are required for RNH* to generate sufficient force to release a SecM stall and indicate that FPA is more nuanced than simply reporting on folding to the native state.

The correlation between the three-state and two-state folding of variants in standard refolding experiments and their ability to read through the SecM stall implies that the intermediate observed in CD and HDX studies off the ribosome is likely populated on the ribosome and again suggests that the general folding trajectory, or pathway, of RNH* is unaltered on the ribosome. These data, together with the pulse proteolysis results, agree with studies of the folding mechanisms of other small proteins on the ribosome, which have shown that the folding trajectory does not change for RNCs relative to free proteins (5, 11). Combining this pulse proteolysis approach with other site-specific mutations in a ϕ-value analysis will further elucidate the folding pathway of ribosome-tethered RNH* I53D and could be applied to other nascent chains to characterize the effect of the ribosome on a range of folding mechanisms.

Our results suggest that the transient folding intermediate of RNase H is responsible for generating the tension required to release the SecM stall. This is perhaps not surprising given that the intermediate of RNase H has a stability comparable to many globular proteins and similar to the small zinc-binding peptide used in the development of FPA (21). Notably, the stability of the intermediate is quite low, suggesting that a stability of just one to two kcal/mol is enough to induce release. Perhaps more surprising, however, is that a stable variant of RNH* without this kinetic intermediate is incapable of release.

What properties of the folding of RNH* I53D prevent the formation of force sufficient to cause release? Perhaps the answer lies in the kinetics of folding. The two-state variants of RNH* are known to fold significantly slower than the three-state variants – the same regions that stabilize the kinetic intermediate are involved in the rate-limiting step, or transition state, for folding (33). Previous bulk experiments have shown that RNH* I53D folds at a rate about four times lower than RNH*. From our pulse-proteolysis experiments, we infer an off-ribosome folding rate of 0.1 s^-1^ for RNH* I53D, relative to *k*_fold_ = 0.74 s^-1^ for RNH* off the ribosome in bulk CD experiments (27). These questions might be best answered by simulations, which can dissect the mechanism of force generation (5, 11, 25).

In addition to the specific results we find for RNH*, the approaches used here (equilibrium and kinetic pulse proteolysis, together with FPA) are a start to a detailed quantitative comparison of protein folding on and off the ribosome, which will lead to a better understanding of protein folding in the cell. FPA has already been used to examine the role of chaperones, such as trigger factor, in the folding of nascent chains (26). Future work can look toward pairing these quantitative experiments with structural studies, such as HDX, to elucidate the folding trajectory in complex, cellular-like environments and could eventually be expanded to investigate the influence of the vectorial and kinetic aspects of translation.

## METHODS

### Generation of plasmids for pulse proteolysis

The coding sequence for RNH* I53D was cloned into the DHFR Control Template provided by New England Biolabs with the PURExpress kit via NdeI and KpnI restriction sites. GS linkers and a SecM stalling sequence with an N-terminal extension (EFLPYRQFSTPVWISQAQGIRAGPQ) were added to the C-terminus of the RNH* I53D coding sequence by around-the-horn mutagenesis (20). RNH* D10A/I53D constructs were generated by subsequent around-the-horn mutagenesis. All constructs were verified by sequencing.

### Preparation of samples for pulse proteolysis

Using PURExpress kits (New England Biolabs), 37.5 μL IVT reactions were assembled on ice with 1.5 μl RNase inhibitor, murine (New England Biolabs), 2 µL FluoroTect GreenLys (Promega), and 375 ng of plasmid encoding the protein of interest. Standard PURExpress kits (E6800S) were used to produce off-ribosome samples, and PURExpress ΔRF123 kits (E6850S) were used to generate RNCs in the absence of release factors. IVT reactions were incubated at 37 °C for 45 min to 2 h.

### Pulse proteolysis

To monitor off-ribosome kinetics, we treated the samples with RNase A before carrying out pulse proteolysis. RNase A (Sigma Aldrich) was added to IVT reactions at a final concentration of 1 mg/mL and incubated for 15–30 min at 37 °C. These reactions were diluted to 65 μL in 1x HKMT (25 mM HEPES, pH 7.4, 15 mM Mg(OAc)_2_, 150 mM KCl, 0.1 mM TCEP). For on-ribosome samples, stalled RNCs were pelleted by ultracentrifugation on a 94 μL 1 M sucrose cushion in 1x HKMT for 40 min at 200,000 x g. The pellet was washed twice with 100 μL 1x HKMT and resuspended in 65 μL 1x HKMT. Measurements of stability by pulse proteolysis were performed as previously described (20). To measure unfolding kinetics, urea stock solutions were set up such that dilution with 49.5 μL of either free protein or RNCs would result in the desired final urea concentration in 1× HKMT. The sample to be tested was rapidly diluted into the urea stock, pipetted up and down to mix, and for each time point, 10 μL was removed and pulsed into a tube containing 1 μL of 2 mg/mL thermolysin (Sigma Aldrich) for 1 min before quenching with 3 μL 500 mM EDTA, pH 8.5.

On-ribosome samples were incubated in 1 mg/mL RNase A after pulse proteolysis for 15–30 min at 37 °C. Samples were mixed with SDS-PAGE loading dye, loaded onto 4-12% or 12% NuPAGE Bis-Tris gels (ThermoFisher), and run in MES buffer for 50 min at 170 V at 4 °C. A full-length RNH* marker was run to aid in quantification. Gels were imaged on a Typhoon FLA9500 (GE Healthcare) with a 488 nm laser and a 510LP filter. Analysis of band intensities was performed as described previously using ImageJ (20). All urea concentrations were determined after the dilution with RNCs or released protein by refractometer, as previously (18).

### CD stability and unfolding kinetics

The plasmid for RNH* I53D was constructed previously (27) and was expressed and purified from inclusion bodies as previously described (34). Data were recorded on an Aviv 430 CD spectropolarimeter at 225 nm and 25 °C in a 0.5-cm pathlength cuvette in 1x HKMT buffer. All urea concentrations were determined by refractometer. Equilibrium denaturation experiments were performed after incubating protein samples overnight at the appropriate urea concentrations. Samples were stirred for 60 s prior to measurement, and data was recorded for 60 s and averaged for each urea concentration. Unfolding kinetics were initiated by rapidly mixing a 100 μM stock of RNH* I53D at 0.2 M urea 1:3 (v/v) in 1x HKMT and the appropriate urea concentration to reach a final denaturant concentration of 4-8 M urea. Samples were manually mixed and dead-times ranged from 15-20 s. The dead-time was added to the start of each trace before fitting. All data were plotted and fit using MatLab as described previously (27).

### Cloning for force-profile constructs

The RNH* coding sequence was inserted into a pRSETA plasmid containing fragments of LepB and a SecM stalling sequence under the control of a T7 promoter as described previously (35). This created a set of pRSETA plasmids containing RNH* with 20 different linker lengths from the RNH* C-terminus to the SecM stall sequence. Linker lengths range from 21 to 61 residues. Variants of RNH* were made by around-the-horn mutagenesis on this set of 20 plasmids. All constructs were verified by sequencing.

### Force-profile analysis

RNCs tethered with varying linker lengths will range in the tension induced on the SecM stall sequence during folding. A very short linker will inhibit protein folding due to occlusion of a portion of the protein in the exit tunnel or interactions between the ribosome and the nascent chain. Extending the linker will allow the protein to fold, and the tension generated will be proportional to the fraction of nascent chains that release the SecM stalling sequence (*f*_FL_). At longer linker lengths, protein folding is distant from the PTC, and there is no coupling between folding and the stall sequence.

RNCs were generated using standard PURExpress kits (New England Biolabs). A 240 μL IVT master mix was assembled on ice, containing 10 μl RNase inhibitor, murine (New England Biolabs) and 12 μL of either EasyTag EXPRESS^35^S Protein Labeling Mix (Perkin Elmer) or L-[^35^S]-Met (Perkin Elmer). For each sample, 9 μL of this master mix was added to 1 μL of plasmid DNA (∼250 ng/μL) and incubated at 37 °C for exactly 15 min before quenching with 1 μL 10 mg/mL RNase A and 1 μL 20 mM chloramphenicol. Samples were incubated at 37 °C for an additional 15 min, mixed with 3 μL 4x LDS loading dye (New England Biolabs) and loaded onto 4-12% NuPAGE Bis-Tris gels (ThermoFisher). The gels were run in MES buffer for 75 min at 150 V at room temperature.

### Analysis of force-profile data

Gel band intensities were determined by plotting the signal intensity for a cross-section of each lane in ImageJ. Data were fit to a bimodal Gaussian distribution in MatLab to calculate *f*_FL_ using the equations

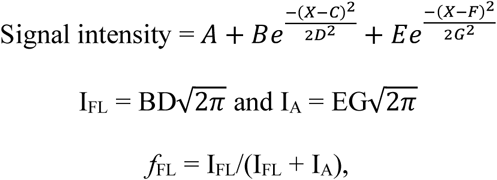

where I_FL_ is the intensity of the full-length band and I_A_ is the intensity of the stalled band.

The following *f*_FL_ were calculated as the average of two experiments: WT L23, L27, L31, L33, L37, L39, L43, L45; I53V L21, L23, L27, L29; I53F L21, L23, L27, L29, L31, L33, L35, L37, L39, L41; NF L43. NF L23, L27, L37, L39, and L47 are the average of three experiments. All other *f*_FL_ shown are the result of a single experiment. The experiment is robust and has been shown to be highly reproducible (5, 22).

## ACKNOWLEDGEMENTS

We would like to thank Brendan Maguire for cloning ∼150 variants of RNH* and different linker lengths for the force-profile work and for helping to purify proteins for CD. We also thank Liang Ming Wee, Lisa Alexander, and Guillermo Chacaltana for their guidance and use of their workspace for the force-profile experiments and the members of the Marqusee Lab for their insights and feedback. We are grateful to Robert Best and Pengfei Tian for their work on simulations and fruitful discussions. We acknowledge funding from the NIH under Grant No. R01-GM050945 (SM) and the NSF GRFP under Grant No. DGE1106400 (MKJ). SM is a Chan Zuckerberg Biohub Investigator. This work is also supported by a grant from the Wellcome Trust (WT095195). JC is a Wellcome Trust Senior Research Fellow. The content is solely the responsibility of the authors and does not necessarily represent the official views of the National Institutes of Health.

## REFERENCES

1. Dobson, C. M. (1999) Protein misfolding, evolution and disease. Trends Biochem. Sci. 24, 329–332

2. Hipp, M. S., Park, S. H., and Hartl, U. U. (2014) Proteostasis impairment in protein-misfolding and -aggregation diseases. Trends Cell Biol. 24, 506–514

3. Kaiser, C. M., Goldman, D. H., Chodera, J. D., Tinoco, I., and Bustamante, C. (2011) The ribosome modulates nascent protein folding. Science. 334, 1723–7

4. Waudby, C. A., Wlodarski, T., Karyadi, M.-E., Cassaignau, A. M. E., Chan, S. H. S., Wentink, A. S., Schmidt-Engler, J. M., Camilloni, C., Vendruscolo, M., Cabrita, L. D., and Christodoulou, J. (2018) Systematic mapping of free energy landscapes of a growing filamin domain during biosynthesis. Biophys. Comput. Biol. 115, 9744–9749

5. Tian, P., Steward, A., Kudva, R., Su, T., Shilling, P. J., Nickson, A. A., Hollins, J. J., Beckmann, R., von Heijne, G., Clarke, J., and Best, R. B. (2018) Folding pathway of an Ig domain is conserved on and off the ribosome. Proc. Natl. Acad. Sci. 115, E11284–E11293

6. Deckert, A., Waudby, C. A., Wlodarski, T., Wentink, A. S., Wang, X., Kirkpatrick, J. P., Paton, J. F. S., Camilloni, C., Kukic, P., Dobson, C. M., Vendruscolo, M., Cabrita, L. D., and Christodoulou, J. (2016) Structural characterization of the interaction of α-synuclein nascent chains with the ribosomal surface and trigger factor. Proc. Natl. Acad. Sci. 113, 5012–5017

7. Sander, I. M., Chaney, J. L., and Clark, P. L. (2014) Expanding Anfinsen’s principle: contributions of synonymous codon selection to rational protein design. J. Am. Chem. Soc. 136, 858–61

8. Clark, P. L. (2004) Protein folding in the cell: reshaping the folding funnel. Trends Biochem. Sci. 29, 527–34

9. Frydman, J., Erdjument-Bromage, H., Tempst, P., and Hartl, F. U. (1999) Co-translational domain folding as the structural basis for the rapid de novo folding of firefly luciferase. Nat. Struct. Biol. 6, 697–705

10. Samelson, A. J., Bolin, E., Costello, S. M., Sharma, A. K., O’Brien, E. P., and Marqusee, S. (2018) Kinetic and structural comparison of a protein’s cotranslational folding and refolding pathways. Sci. Adv. 4, eaas9098

11. Guinn, E. J., Tian, P., Shin, M., Best, R. B., and Marqusee, S. (2018) A small single-domain protein folds through the same pathway on and off the ribosome. Proc. Natl. Acad. Sci. 115, 12207–12211

12. Ozkan, S. B., Bahar, I., and Dill, K. A. (2001) Transition states and the meaning of Phi-values in protein folding kinetics. Nat. Struct. Biol. 8, 765–769

13. Holtkamp, W., Kokic, G., Jäger, M., Mittelstaet, J., Komar, A. A., and Rodnina, M. V. (2015) Cotranslational protein folding on the ribosome monitored in real time. Science. 350, 1104–1107

14. Mercier, E., and Rodnina, M. V (2018) Co-translational Folding Trajectory of the HemK Helical Domain. Biochemistry. 57, 3460–3464

15. Goldman, D. H., Kaiser, C. M., Righini, M., and Bustamante, C. (2015) Mechanical force releases nascent chain–mediated ribosome arrest in vitro and in vivo. Science (80-.). 348, 457–460

16. Liu, K., Rehfus, J. E., Mattson, E., and Kaiser, C. M. (2017) The ribosome destabilizes native and non-native structures in a nascent multidomain protein. Protein Sci. 10.1002/pro.3189

17. Park, C., and Marqusee, S. (2006) Quantitative determination of protein stability and ligand binding by pulse proteolysis. Curr. Protoc. Protein Sci. 46, 20.11.1-20.11.14

18. Park, C., and Marqusee, S. (2005) Pulse proteolysis: A simple method for quantitative determination of protein stability and ligand binding. Nat. Methods. 2, 207–212

19. Na, Y.-R., and Park, C. (2009) Investigating protein unfolding kinetics by pulse proteolysis. Protein Sci. 18, 268–276

20. Samelson, A. J., Jensen, M. K., Soto, R. A., Cate, J. H. D., and Marqusee, S. (2016) Quantitative determination of ribosome nascent chain stability. Proc. Natl. Acad. Sci. 113, 13402–13407

21. Nilsson, O. B., Hedman, R., Marino, J., Wickles, S., Bischoff, L., Johansson, M., Müller-Lucks, A., Trovato, F., Puglisi, J. D., O’Brien, E. P., Beckmann, R., and von Heijne, G. (2015) Cotranslational Protein Folding inside the Ribosome Exit Tunnel. Cell Rep. 12, 1533–1540

22. Nilsson, O. B., Nickson, A. A., Hollins, J. J., Wickles, S., Steward, A., Beckmann, R., von Heijne, G., and Clarke, J. (2017) Cotranslational folding of spectrin domains via partially structured states. Nat. Struct. Mol. Biol. 24, 221–225

23. Kemp, G., Kudva, R., de la Rosa, A., and von Heijne, G. (2019) Force-Profile Analysis of the Cotranslational Folding of HemK and Filamin Domains: Comparison of Biochemical and Biophysical Folding Assays. J. Mol. Biol. 431, 1308–1314

24. Farías-Rico, J. A., Ruud Selin, F., Myronidi, I., Frühauf, M., and von Heijne, G. (2018) Effects of protein size, thermodynamic stability, and net charge on cotranslational folding on the ribosome. Proc. Natl. Acad. Sci. U. S. A. 115, E9280–E9287

25. Leininger, S. E., Trovato, F., Nissley, D. A., and O’Brien, E. P. (2019) Domain topology, stability, and translation speed determine mechanical force generation on the ribosome. Proc. Natl. Acad. Sci. 116, 5523–5532

26. Nilsson, O. B., Müller-Lucks, A., Kramer, G., Bukau, B., and Von Heijne, G. (2016) Trigger Factor Reduces the Force Exerted on the Nascent Chain by a Cotranslationally Folding Protein. J. Mol. Biol. 428, 1356–1364

27. Spudich, G. M., Miller, E. J., and Marqusee, S. (2004) Destabilization of the Escherichia coli RNase H Kinetic Intermediate: Switching Between a Two-state and Three-state Folding Mechanism. J. Mol. Biol. 335, 609–618

28. Rosen, L. E., Kathuria, S. V., Matthews, C. R., Bilsel, O., and Marqusee, S. (2015) Non-Native Structure Appears in Microseconds during the Folding of E. coli RNase H. J. Mol. Biol. 427, 443–453

29. Connell, K. B., Horner, G. A., and Marqusee, S. (2009) A Single Mutation at Residue 25 Populates the Folding Intermediate of E. coli RNase H and Reveals a Highly Dynamic Partially Folded Ensemble. J. Mol. Biol. 391, 461–470

30. Connell, K. B., Miller, E. J., and Marqusee, S. (2009) The Folding Trajectory of RNase H Is Dominated by Its Topology and Not Local Stability: A Protein Engineering Study of Variants that Fold via Two-State and Three-State Mechanisms. J. Mol. Biol. 391, 450–460

31. Cecconi, C., Shank, E. A., Bustamante, C., and Marqusee, S. (2005) Direct observation of the three-state of a single protein molecule. Science. 4174, 2057–2060

32. Raschke, T. M., Kho, J., and Marqusee, S. (1999) Confirmation of the hierarchical folding of RNase H: a protein engineering study. Nat. Struct. Biol. 6, 825–831

33. Hu, W., Walters, B. T., Kan, Z., Mayne, L., Rosen, L. E., Marqusee, S., and Englander, S. W. (2013) Stepwise protein folding at near amino acid resolution by hydrogen exchange and mass spectrometry. Proc. Natl. Acad. Sci. U. S. A. 110, 7684–9

34. Rosen, L. E., Connell, K. B., and Marqusee, S. (2014) Evidence for close side-chain packing in an early protein folding intermediate previously assumed to be a molten globule. Proc. Natl. Acad. Sci. 111, 14746–14751

35. Ismail, N., Hedman, R., Schiller, N., and Von Heijne, G. (2012) A biphasic pulling force acts on transmembrane helices during translocon-mediated membrane integration. Nat. Struct. Mol. Biol. 19, 1018–1023

